# Excessive drinking and checking in the rat model of Schedule-Induced Polydipsia reveal impaired bi-directional plasticity at BNST GABA synapses

**DOI:** 10.1101/799452

**Authors:** Staci Angelis, James Gardner Gregory, Emily R. Hawken, Éric C. Dumont

## Abstract

Compulsions, defined by debilitating repetitive actions, permeate many mental illnesses and are challenging to treat partly because of a limited understanding of their neurobiological underpinnings. Accumulating evidence suggests the rodent model of Schedule-Induced Polydipsia (SIP) as a promising pre-clinical assay to elucidate the neurobiological and behavioural manifestations of compulsivity. In the rodent SIP paradigm, susceptible rats develop adjunctive excessive drinking when they are chronically food restricted and presented with food pellets according to a fixed-time schedule. We found that normally, bi-directional plasticity of GABA synapses in the oval bed nucleus of the stria terminalis (ovBNST) tightly followed the rats’ satiety state where low-frequency stimulation-induced potentiation (LTPGABA) prevailed in sated rats whilst food restriction uncovered long-term depression (LTDGABA). In rats that developed excessive drinking during SIP, removing the caloric restriction failed at reverting LTDGABA to LTPGABA whereas bi-directional plasticity at ovBNST GABA synapses was unaltered in low-drinking SIP-trained rats. Excessive drinking ceased in polydipsic rats removed from their caloric restriction; however, these rats retained a form of compulsive schedule-induced checking (SIC) and impaired plasticity at ovBNST GABA synapses for several days following termination of the caloric restriction. We conclude that altered bi-directional plasticity at ovBNST GABA synapses is a neurophysiological trace of compulsivity in susceptible rats in the SIP model.

## Introduction

Compulsions are time-consuming, repetitive actions that cause significant distress and impairments in both the function and quality of daily life (American Psychiatric Association, 2013). Compulsive behaviours are symptomatic of many psychiatric disorders including obsessive-compulsive disorder (OCD), addictions, and schizophrenia (Berman et al., 1995; Drubach, 2015; Everitt and Robbins, 2005; Hariprasad et al., 1980). Despite common etiological characteristics across psychiatric diagnoses, the underlying neurophysiology of compulsivity remains largely unknown. Existing preclinical rodent models may help elucidate where and how in the brain is pathological compulsivity represented (Everitt and Robbins, 2016; Szechtman et al., 2017).

In the schedule-induced polydipsia (SIP) paradigm, food-restricted rats develop adjunctive excessive drinking when presented with food pellets according to a fixed-time schedule (Falk, 1966; Platt et al., 2008). Interestingly, only a subset of rats within any given cohort develops excessive drinking suggesting a powerful pre-clinical tool to understand how environment and genetics interact to trigger the maladaptive behaviour (Moreno and Flores, 2012; Rosenzweig-Lipson et al., 2007; Toscano et al., 2008). Although the drinking behaviour in SIP is clearly excessive, non-physiological, and maladaptive due to the risk of both hyponatremia and death, whether it is truly compulsive remains to be definitively determined (Hawken et al., 2013b). Chronic caloric restriction is a necessary condition to induce and maintain excessive drinking in the SIP paradigm, increasing the salience of scheduled food pellet release (Falk, 1961, 1967, 1971; Hooks et al., 1994). Determining whether food restriction is necessary to maintain adjunctive behaviours of polydipsic rats may further confirm compulsivity in SIP and pave the way to a better understanding of its neurophysiological underpinnings.

In humans, compulsive behaviours may serve to alleviate the symptoms of a chronic stress state (American Psychiatric Association, 2013). In the SIP paradigm, food restriction serves as the chronic homeostatic challenge. Adrenalectomy attenuates excessive drinking, which is subsequently restored by corticosterone replacement, cementing the role of stress in this adjunctive behaviour (Levine and Levine, 1989). Furthermore, access to the drinking spout during SIP decreases plasma corticosterone, and there is an inverse relationship between plasma corticosterone and water intake during SIP (Brett and Levine, 1979, 1981; Mittleman et al., 1988; Tazi et al., 1986). Consequently, brain regions important for energy metabolism should play a key role in converting caloric deficit stress into motivated behaviours in order to re-establish homeostasis.

The Bed nucleus of the Stria Terminalis (BNST) -and in particular its oval (ov) sub-region-plays such an adaptive role but also contributes to the manifestation of maladaptive compulsive-like behaviours in both laboratory animals and humans (Jennings et al., 2013; Krawczyk et al., 2013; van Kuyck et al., 2008; Welkenhuysen et al., 2013). Notably, BNST neurons firing rate increases in SIP rats and high-frequency electrical stimulation of these neurons reduces excessive drinking in SIP (van Kuyck et al., 2008; Welkenhuysen et al., 2013). Likewise, deep brain stimulation in the BNST produces long-lasting reductions in obsessions and compulsions in OCD patients (Luyten et al., 2016).

Here, using brain slice electrophysiology, we saw that satiety state-dependent bi-directional plasticity at ovBNST GABA synapses was specifically impaired in rats developing excessive drinking and checking in the SIP paradigm, revealing a potential neurophysiological trace of compulsivity in the brain.

## Materials and Methods

### Animals

Seventy-two male Long Evans rats weighing 250-275 g (Charles River, St-Constant, QC) were pair-housed in clear Plexiglas cages (45×23×20cm) lined with bedding (Beta Chip, NEPCO, Warrenburg, NY). The cages were located in a climate-controlled colony room (21±1°C; humidity 40-70%) on a reversed light/dark cycle (lights OFF: 8:00-20:00). After a 7-day acclimatization period with *ad libitum* access to food (rat chow; LabDiet rodent feed #5001, PMI Nutrition International, Brentwood, MO) and water, the rats were individually housed for the rest of the study. All the experiments were conducted in accordance with the Canadian Council on Animal Care guidelines for use of animals in experiments and approved by the Queen’s University Animal Care Committee (protocol # 2014-1537).

### Experimental Design

The rats were randomly assigned to six conditions: Free-fed (n=6), Schedule-Induced Polydipsia (SIP, n=28), Chronic Food Restriction (cFDR, n=7), Chronic Food Restriction with acute refeed (cFDR_refeed_, n=9), SIP with acute refeed (SIP_refeed_, n=17) and, SIP with chronic refeed (SIP_c/refeed_, n=5). All the rats had *ad libitum* access to water in their home cage. Free-fed rats were individually housed for 29 consecutive days with *ad libitum* food access. SIP and SIP_refeed_ were restricted to a 1− or 2-hour free-feeding periods (12:00–13:00 or 14:00) for 28 consecutive days immediately following the SIP sessions. We targeted 1 hour free-feeding to counterbalance the 120 food pellets received during the daily SIP sessions. However, we adapted SIP free-feeding time to ensure the rats remained within 85-90% of their pre-food restriction weight. SIP_refeed_ were acutely refed immediately following their last SIP session, on day 28 (14:00-8:00). cFDRs were restricted to a 2-hour free-feeding period (12:00–14:00) for 29 consecutive days. cFDR_refeed_ were restricted to a 2-hour free-feeding period (12:00–14:00) for 28 consecutive days before being acutely refed on day 28 (14:00-8:00)

### SIP paradigm

We used 6 commercial (Med Associates, ST-Albans, VT) operant conditioning chambers placed in ventilated sound-attenuating cabinets. A food dispenser was located opposite to a metal, ball-bearing, drinking spout within each chamber. The operant conditioning chambers were controlled and data acquired by a computer running MED-PC-IV (Med Associates Inc., St. Albans, VT). After 7 days of acclimatization to the food restriction regime, 2 SIP groups (SIP, SIP_refeed_) had daily (2 hours) access to the operant conditioning chambers for a minimum of 21 consecutive days at approximately 10:00. During those 2 hours, 45mg dustless precision food pellet (Bio-Serv, Frenchtown, NJ) were delivered in the food dispenser trays on a fixed-time 60-sec schedule. The rats had unlimited access to the water spout during the SIP session and intake was determined by weighing the bottles before and after each session. SIP,SIP_refeed_, SIP _c/refeed_ rats were classified as High Drinkers (SIP-HD, SIP-HD_refeed_, SIP-HD_c/refeed_, respectivly if they drank 15mls for 3 consecutive days) (Gregory et al., 2015; Hawken and Beninger, 2014; Hawken et al., 2013a; Hawken et al., 2011; Hawken et al., 2013b). Otherwise, rats were classified as Low Drinkers (SIP-LD, SIP-LD_refeed_ or SIP-LD_C/Refeed_). Rats in the SIP_c/refeed_ were chronically food restricted for 7 days and acutely refed (22 hours) before their first SIP session to obtain a sated-state baseline water intake value. The rats were returned to their food restricted regime immediately following this first SIP session and daily SIP training resumed for at least 21 consecutive days. SIP_c/refeed_ rats were then removed from food restriction after meeting criteria for HD for at least 10 consecutive days. SIP training continued daily while rats were fed *ad libitum* until operant chamber water intake decreased significantly below values of the 21_st_ SIP session; this took between 3 and 8 days. Head-entry duration at the water-spout (ms) was recorded in 5-sec bins over the 1-min interval between food pellet deliveries for all SIP_c/refeed_ to quantify water spout checking. The first 30 seconds following food pellet delivery was considered adjunctive and the final 30 seconds was considered non-adjunctive (Lopez-Crespo et al, 2004).

### Brain Slice Preparation and Electrophysiology

The rats were euthanized (approximately 10:00) under deep isoflurane anesthesia (5% at 5 L/min). The brains were rapidly removed and kept in an iced-cold physiological solution containing (in mM): 126 NaCl, 2.5 KCl, 1.2 MgCl_2_, 6 CaCl_2_, 1.2 NaH_2_PO_4_, 25 NaHCO_3_ and 12.5 D-glucose, equilibrated with 95%O_2_/5%CO_2_. The brains were cut in coronal slices (250 μm) with a vibrating-blade microtome (Leica VT-1000, Leica Canada, Concord, ON) in the physiological solution (2°C). Brain slices containing the BNST were incubated at 34°C for 60 mins before being transferred to a recording chamber constantly perfused (3 mls/min) with the physiological solution (34°C). Slices not immediately used following the 60 mins incubation were kept in the physiological solution at room temperature until further use. GABAA-IPSCs were recorded in the whole-cell voltage-clamp configuration using glass micropipettes (3.5 MOhm) filled with a solution containing (in mM): 70 Cs+MeSO_3−_, 58 KCl, 0.5 EGTA, 7.5 HEPES, 1.2 MgCl_2_, 12 NaCl, 1 Mg-ATP, 0.3 GTP, and 1 phosphocreatine. ECl in these conditions was approximately −32mV such that evoked GABA_A_ postsynaptic responses were inward currents when neurons were voltage-clamped at −70mV. Detailed methodology for recordings of ovBNST GABAA-IPSC was previously published (Krawczyk et al., 2011). GABAA-IPSCs were evoked by local fiber stimulations with tungsten electrodes (FHC, Bowdoin, ME) using a bipolar stimulus isolator (World Precision Instruments, Sarasoto, FL) in the presence of the AMPA antagonist DNQX (50μM). Electrodes were placed in the ovBNST, 100-500 μm dorsal from the recorded neurons, and GABAA-IPSCs we evoked (10-100μA, 0.1ms duration) at 0.1Hz. Following a 5-min steady baseline recording, neurons were subjected to a low-frequency stimulation (LFS) synaptic plasticity-inducing protocol (1Hz, 5 mins), followed by a 25-min (minimum) post-induction recording. We quantified total charge transfer and defined 3 possible outcomes for the LFS protocol: 1-LFS-induced long-term potentiation (LTPGABA; >20% deviation from baseline), 2-LFS-induced long-term depression (LTDGABA; <20% deviation from baseline) or 3-no change (NC), 20 mins post-LFS. Every recording was verified *a posteriori* and any indication that a change in IPSC magnitude or kinetic correlated with a change in recording quality (1mv, 5ms test pulse) resulted in exclusion.

### Drugs

Stock solution of 6,7-dinitroquinoxaline-2,3-dione (DNQX; 100mM; RandD Systems, Minneapolis, MN) was prepared in dimethyl sulfoxide (DMSO; 100%; Fisher Canada, Ottawa, ON) then further dissolved in the physiological solution at the desired concentration (final DMSO concentration ≤0.1%).

### Statistical Analyses

We used one- and two-ways ANOVAs to compare variance between groups and conducted tests of simple effects when ANOVAs yielded significance. To control for type I error across multiple comparisons, a Bonferroni correction was applied to all p-values, unless otherwise stated. In order to investigate changes in GABAA-IPSCs total charge transfer following LFS, the baseline area under the curve (AUC) was compared to the post-LFS AUC [(post AUC – baseline AUC)/ baseline AUC) * 100]. Measurements of the AUC was done with Axograph X. In graphs denoting electrophysiology time-course, each data point represents the average of 1-min bins (6 evoked GABAA-IPSCs) across recorded neurons. Neuronal responses were compared using the Fisher’s exact probability test across all groups. Finally, the relation between water consumed on final SIP acquisition session and 20 min post-LFS GABAA-IPSCs was analyzed using a Pearson’s correlation coefficient. *P values* ≤ 0.05 were considered statistically significant. Data are reported as mean ± SEM. All statistical analyses were done with SPSS 24 (SAS Institute).

## Results

We combined the rodent model of SIP and brain slice electrophysiology to identify a potential neurophysiological trace (s) of compulsivity. We specifically compared bi-directional plasticity at ovBNST GABA synapses between several groups of rats including high (HD) and low (LD) drinkers in the SIP paradigm. We refed a group of established SIP-HD rats to remove the metabolic challenge required to induce excessive drinking in order to detect potential compulsive behavioural components and neurophysiological traces in the SIP paradigm.

### Excessive drinking in sub-populations of male rats undergoing the SIP paradigm

As expected from the established experimental SIP protocol, a sub-group of rats gradually increased their water consumption in the operant chambers over the course of the daily 2-hour SIP sessions (Two-way mixed ANOVA, main effect of time (days); F_20,560_=17.6, *p*≤0.001, Fig. 1B). Those rats reached criterion for excessive drinking in 11±1 days and were classified as HD (Fig. 1B). Although all the rats gradually reduced their daily home cage drinking, HD progressively and excessively drank during the 2-hour SIP sessions (Fig. 1C; *F*_20, 560_= 17.66, *p*≤ 0.001). In contrast, rats classified as LD retained a stable total daily water intake throughout SIP training, reducing home cage drinking whilst increasing operant chamber drinking by a similar amount (Fig. 1C)

**Figure 1.**
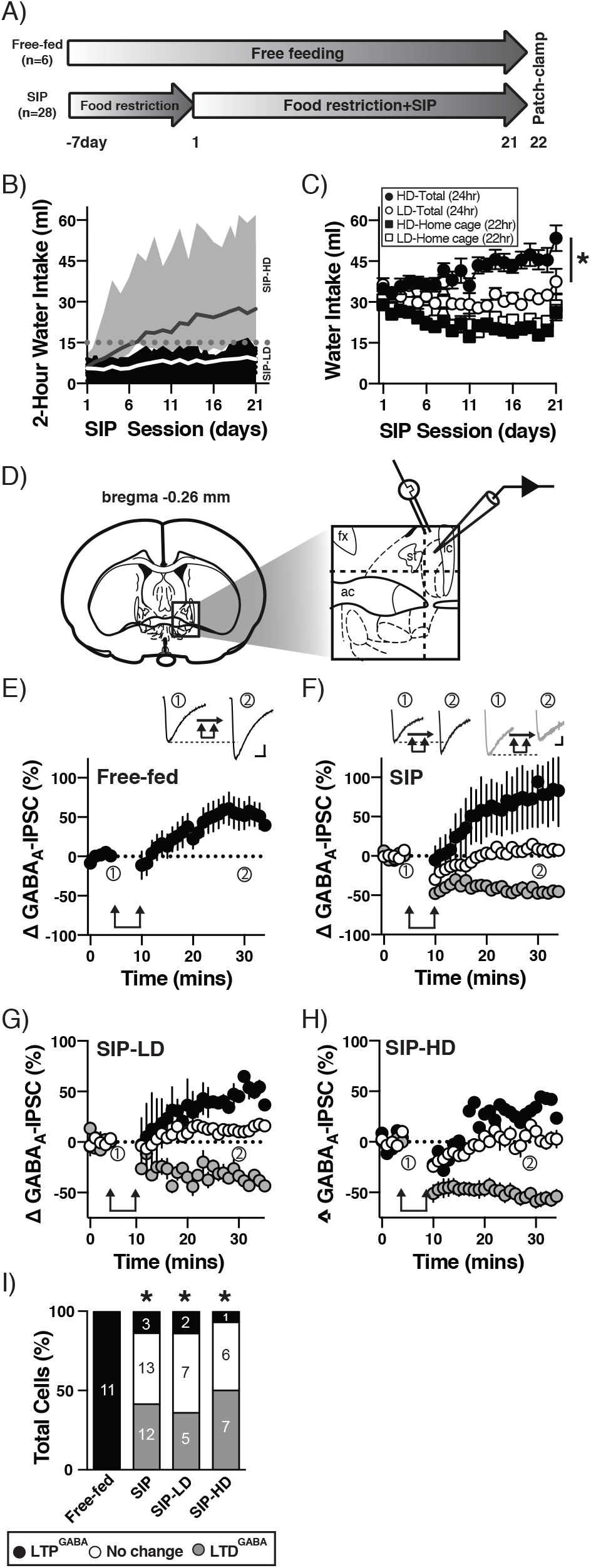
Effect of the SIP behavioural paradigm on bi-directional plasticity at ovBNST GABA synapses. A, Experimental timeline for rats included in the Free-fed and SIP groups. B, Two-hour operant chamber water intake in Low Drinkers (LD; white line, black shaded area) and High Drinkers (HD; black line, grey shaded area). Shaded areas represent the range of water intake across individual rats. C, Total (home-cage + operant chamber) or home-cage water intake as a function of daily SIP sessions. * *p*<0.05 main effect of group comparing LD-Total (24hr) and HD-Total (24hr) over the 21 sip sessions. D, Schematic illustrating stimulating and recording electrodes placements within the ovBNST (adapted from Paxinos and Watson, 2005). Dotted lines indicate the medial and ventral limits of the recordings that were accordingly restrained to the oval (ov) BNST. E-H, Binned (1 mins, 6 events) electrically-evoked ovBNST GABAA-IPSCs as a function of time recorded in brain slices prepared from (E) Free-fed (n_cells_= 11; n_rats_=6), (F) SIP (n_cells_= 29; n_rats_=14), (G) SIP-LD (n_cells_= 14; n_rats_=7), (H) SIP-HD (n_cells_= 17; n_rats_=7), Insets in E and F are representative GABAA-IPSCs before and 20 mins after LFS (represented by vertical arrows symbol). Black traces in E inset show a representative LFS-induced LTPGABA. Grey traces in F inset show a representative LFS-induced LTDGABA. Scale bars: 200pA, 10ms. I, Percentage of cells responding to LFS as either LTPGABA, No Change or LTDGABA across experimental conditions. Circled numbers in panels E and F indicate the time-intervall used to generate the representative traces. * *p*<0.05 compared to Free-fed.

### Impaired bi-directional plasticity at ovBNST GABA synapses with excessive drinking

Electrical local fiber stimulation at 0.1Hz in the presence of the AMPA antagonist DNQX (50μM) evoked reliable whole-cell GABAA-IPSCs in the ovBNST. Following a minimum of 5 mins of stable GABAA-IPSCs, LFS (1 Hz, 5 mins) resulted in robust LTPGABA in all ovBNST neurons recorded from Free-fed rats (Fig. 1E, I). SIP training revealed that LFS-induced GABA plasticity in the ovBNST was bi-directional, uncovering both LTPGABA and LTDGABA, in addition to no net change in GABA-IPSC magnitude in 41, 45, and 14% of recorded neurons, respectively (Fig. 1F, I; *p*≤ 0.001). SIP-induced bi-directional plasticity of ovBNST GABA synapses occurred regardless of behavioural outcome, with comparable percentages of LTPGABA, LTDGABA, and no changes in both LD and HD (Fig. 1G, H, I; LD, *p*≤ 0.001; HD, *p*≤ 0.001, compared to Free-fed).

### Food restriction and refeed reveal bi-directional plasticity at ovBNST GABA synapse

Food restriction was sufficient to uncover bi-directional plasticity of ovBNST GABA synapses (Fig. 2B). In slices prepared from cFDR rats, LFS resulted in all 3 possible outcomes: LTPGABA, LTDGABA, or no change in 25, 31, and 44% of neurons, respectively. Acutely (18 hours) refeeding food restricted rats (cFDR_Refeed_) after 28 days of food restriction had little effect on bi-directional ovBNST GABA plasticity with LTPGABA, LTDGABA, or no change in 42, 29, and 29% of neurons, respectively (Fig. 2C, F). However, refeeding SIP rats completely abolished LTDGABA in low drinkers (SIP-LD_Refeed_, Fig. 2D, F) but not in high drinkers (SIP-HD_Refeed_ Fig. 2E, F), suggesting a possible link between excessive drinking and the inability of ovBNST GABA synapses to detect reestablishment of metabolic state. Consequently, the 2-hour water intake of SIP_Refeed_ rats negatively correlated with the magnitude of LFS-induced LTPGABA, further confirming refeed-resistant LTDGABA in the highest drinkers (r =−0.6, *p=*0.005, Fig. 2G).

**Figure 2.**
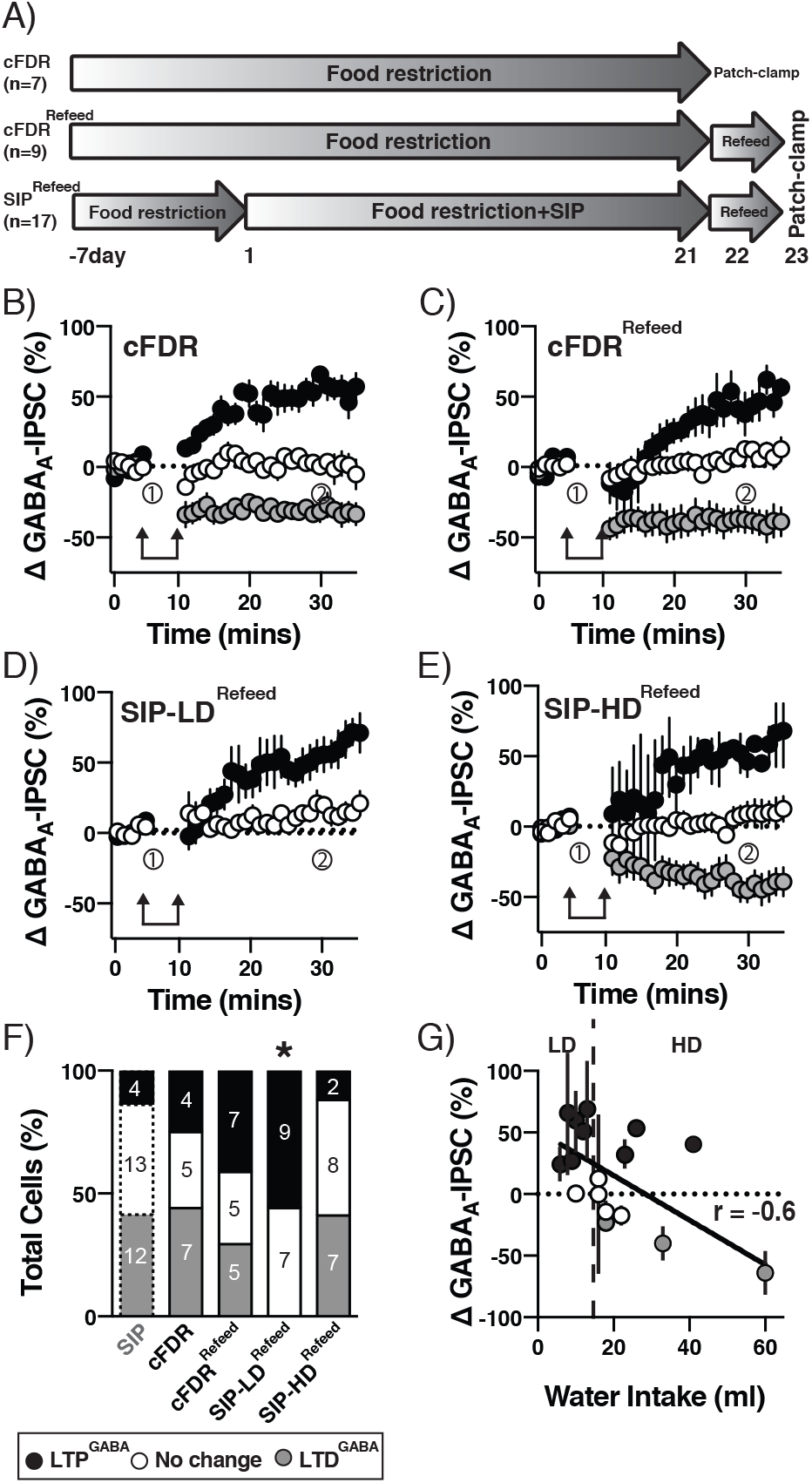
Effect of chronic food restriction, SIP, and acute 18 hour refeeding on bi-directional plasticity at ovBNST GABA synapses. A, Experimental timeline. cFDR_refeed_, SIP-LD_refeed_ and SIP-HD_refeed_ rats were refed for 18 hours. B-E, Binned (1 mins, 6 events) electrically-evoked ovBNST GABAA-IPSCs as a function of time recorded in brain slices prepared from (B) cFDR (n_cells_= 16; n_rats_=7), (C) cFDR-refeed (n_cells_= 17; n_rats_=9), (D) SIP-LD_refeed_ (n_cells_= 16; n_rats_=10), and (E) SIP-HD_refeed_ (n_cells_= 17; n_rats_=7). F, Percentage of cells that responded to LFS as either LTPGABA, No Change or LTDGABA across experimental conditions. * *p*<0.05 compared to cFDR. G, GABAA-IPSC magnitude at 20 mins post-LFS as a function of water intake on the last SIP training day in SIP-HD_Refeed_ and SIP-LD_Refeed_ rats (r =−0.6, *p* = 0.005). Each data point is the average GABAA-IPSCs from all neurons for each individual rat.

### Extinguishing excessive drinking revealed adjunctive Schedule-induced Checking (SIC) at the water spout

Five HD rats (SIP_c/refeed_) were returned to *ad libitum* access to rodent chow after several days of SIP training (Fig. 3A). Upon *ad libitum* access to food, 4 of 5 SIP_c/refeed_ rats rapidly reduced their drinking back to SIP day 1 values within 3 to 8 days during the daily 2-hour SIP sessions (Final SIP session vs. 1_st_ SIP session, t_4_=−2.3, *p*=0.2, Fig. 3B). One SIP_c/refeed_ drinking rat was halved but continued to meet criteria for HD after 8 days of *ad libitum* access to rodent chow. Although the SIP_c/refeed_ rats reduced their water consumption upon termination of the food restriction regime, they continued to check the water spout (Fig. 3C). Checking was adjunctive since it was restricted to 30 secs immediately following food pellet delivery (within-subject effects of SIP session for the 0-30 secs, *F*_3,12_=10.6, *p*≤ 0.001 and 31-60 secs, *F*_3,12_=1.7, *p*=0.2, Fig. 3D). Whole-cell recordings in slices prepared from those HD rats that had multiple days of *ad libitum* access to food (SIP_c/refeed_) and demonstrating SIC revealed that bi-directional GABA plasticity did not revert to Free-fed-like conditions with LTPGABA, LTDGABA, or no change in 19, 50, and 31% of neurons, respectively (Fig. 3E).

**Figure 3.**
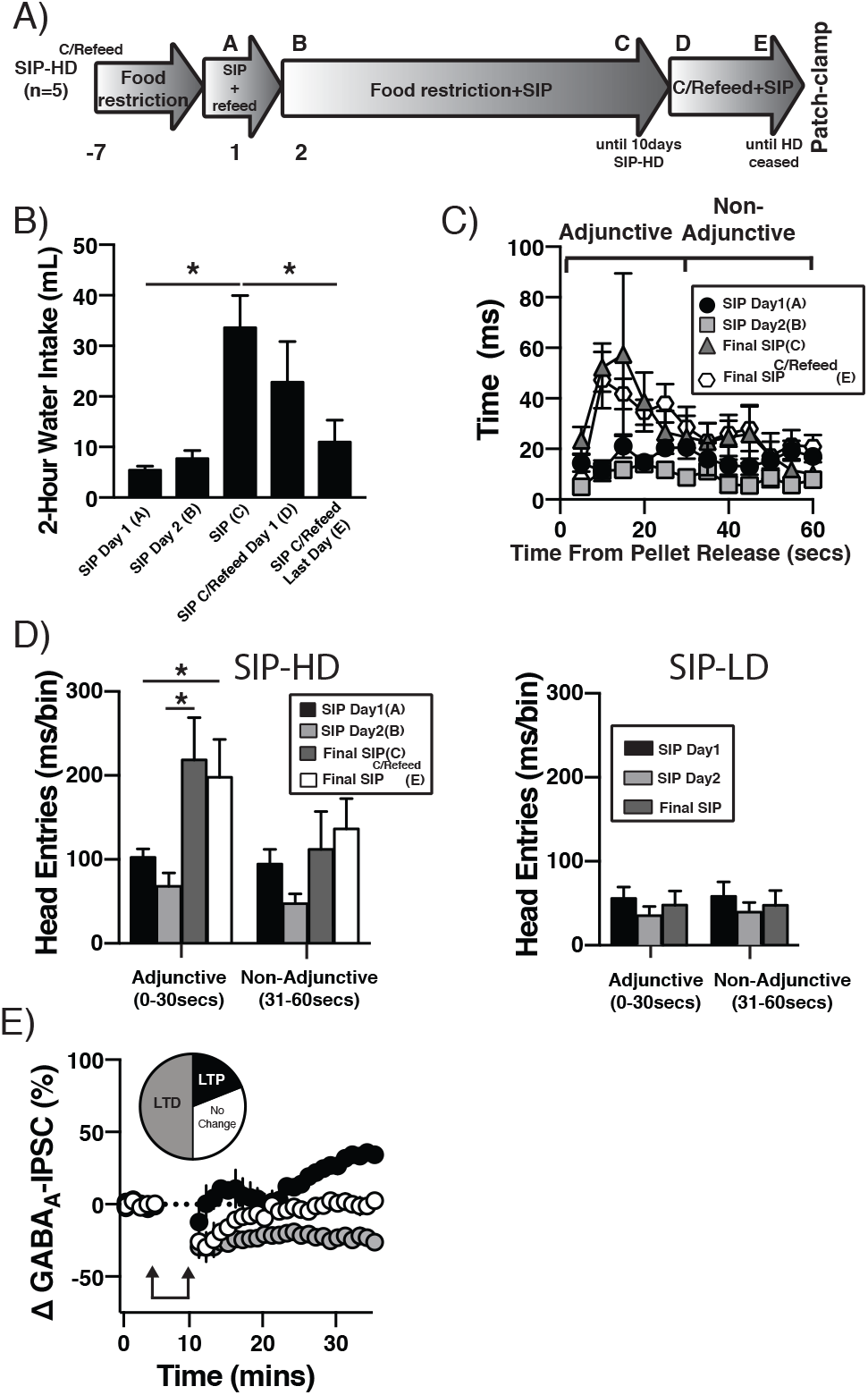
Effect of chronic refeeding of HD-SIP rats on water intake, water spout checking, and bi-directional plasticity at ovBNST GABA synapses. A, Experimental timeline. SIP_c/refeed_ were refed for at minimum 66 hours. B, Bar graph illustrating the effect of SIP training and chronic refeeding on 2-hour operant chamber water intake. C, Head-entry duration at the water-spout recorded in 5-sec bins over the 1-min interval between food pellet deliveries for all SIP_c/refeed_ rats. D, Average head-entry duration during the adjunctive (first 30 secs) and non-adjunctive (last 30 secs) following food pellets presentation at 4 experimental time-points in SIP-HD_C/Refeed_ (left) and SIP-LD_C/Refeed_ (right) rats. E, Binned (1 mins, 6 events) electrically-evoked ovBNST GABAA-IPSCs as a function of time recorded in brain slices prepared from SIP-HD_C/Refeed_ (n_cells_= 16; n_rats_=5). Pie chart shows the percentage of neurons responding to LFS as either LTPGABA (No Change or LTDGABA. * *p*<0.05

## Discussion

Accumulating evidence supports the rodent SIP paradigm as a pre-clinical model to study the neurobiology of compulsivity (Belin-Rauscent et al., 2016; Moreno and Flores, 2012; Platt et al., 2008). Here, by removing the caloric restriction necessary to induce SIP, we saw that although excessive drinking vanished, the rats continued to adjunctively check the water spout compulsively, i.e. in a time-consuming and repetitive way, without any obvious benefits. Supporting this behavioural evidence of compulsivity, we saw that bi-directional plasticity of GABA synapses in the BSNT remained ‘locked’ in its food-restricted mode in HD and SIC rats upon caloric replenishment, but not in those that resisted excessive drinking (LD).

### Locked bi-directional plasticity at ovBNST GABA synapses of HD rats

Here, we hypothesized that excessive drinking behaviour in the rat model of SIP is both compulsive and associated with a neurophysiological signature in the BNST, a brain region associated with compulsive behaviours in laboratory animals and humans (Luyten et al., 2016; Welkenhuysen et al., 2013). In the rodent SIP model, the development of excessive drinking requires long-term caloric restriction which acts to increase the rewarding salience of the food pellets (Lockie and Andrews, 2013). The synergistic interaction between scheduled delivery of food pellets with their enhanced rewarding salience results in excessive drinking in rats, in a subject-specific way (Falk, 1967). Historically, rats have been categorized as polydipsic when water intake during the 2-hour SIP sessions exceeded 22 hours of home cage drinking (Flory, 1971). Recently, we widened this criterion to classify rats as HD if their operant chamber water intake was equal or greater than 50% of home-cage intake (Hawken et al., 2013b). We detected neurophysiological evidence supporting this criteria, showing a clear impairment in satiety-state dependent bi-directional GABA plasticity of ovBNST synapses in SIP-HD_Refeed_ compared with SIP-LD_Refeed_ rats. In all tested SIP-LD_Refeed_ rats, acute refeeding following long-term food restriction eliminated LTDGABA, mostly reverting LFS-induced GABA plasticity to Free-fed sated-state values. In contrast, LFS-induced bi-directional GABA plasticity in the ovBNST of all HD rats fully resisted acute refeeding, i.e., remained dominated by LTDGABA and no change: it did not revert to sated-state LTPGABA.

In brain slices prepared from sated rats, a 5min/1Hz (LFS) plasticity protocol inevitably results in robust LTPGABA in the ovBNST (Gregory et al., 2015). Eighteen hours of fasting uncovers LTDGABA and consequently, plasticity at ovBNST GABA synapses is bi-directional and tightly follows acute satiety states (Gregory et al., 2015). Likewise, when rats were chronically food restricted and acutely refed we herein show that the direction of LFS-induced GABA plasticity was also satiety-state dependent. This was particularly perceptible in the sub-group of SIP-LD rats whereby an acute 18-hour refeed completely eliminated LTDGABA despite 28 days of food restriction and 21 days of SIP training. In contrast, bi-directional plasticity completely resisted 18-hours or several days of refeed in HD rats, that is, GABA synapses remained in their depressed (LTDGABA or no change) mode. Therefore, we can conclude, first, that it is most likely the caloric restriction that uncovered bi-directional plasticity at ovBNST GABA synapses in SIP-trained rats. Second, that caloric intake-dependent bi-directional ovBNST GABA plasticity may predict the behavioural outcome in the SIP paradigm (HD vs. LD). The cellular/molecular mechanism (s) responsible for bi-directional plasticity at these synapses is currently unknown but seems a vulnerability node in rats developing excessive drinking in the SIP paradigm.

### Schedule-induced checking (SIC) in chronically refed HD rats

Food restriction is required to induce excessive drinking in the SIP paradigm. However, it remained unclear whether it is also necessary to maintain the maladaptive behaviour. To test this possibility, we refed HD rats and saw that within 3-8 days of *ad libitum* access to food, excessive drinking in the operant chamber ceased completely in 4 rats and dropped by 50% in a 5_th_ rat. Yet, all the rats continued to adjunctively check the water spout, revealing two distinguishable behaviours once SIP was established. First, excessive water intake which was contingent on caloric restriction and second, a form of SIC that persisted even if the rats were refed for several days. SIC was adjunctive to food pellet delivery, suggesting that although rats were sated, scheduled intermittent access to food retained powerful behavioural triggering effects. Future experiments should determine whether SIC remains adjunctive with a variable schedule of food pellet delivery. It would also be interesting to determine whether food remains necessary or if secondary food predicting cues could trigger SIC as well. Together, these data could further support the compulsive nature of SIC. In addition, our data show that bi-directional plasticity of ovBNST GABA synapses could not recover from days of free-feeding in all tested SIC rats, suggesting this neurophysiological trace is closely related to this potentially compulsive aspect of the SIP paradigm.

The SIC behaviour we observed meets many of the criteria for compulsive checking in humans and may consequently have face validity of the clinical condition. First, compulsive checking in humans is time-consuming, repetitive and interferes with normal daily life activities (American Psychiatric Association, 2013). In SIC, the water spout became the key location in which the rat spent a significant amount of time without any observable benefit during the 2-hour SIP sessions. Future experiments could determine whether SIC actually significantly narrows the behavioural repertoire of the rats compared with non-SIC rats. Second, compulsive checking in humans is triggered either by an anxiety-producing cognitive disturbance (fear that the stove will set the home on fire), a physical trigger (physically seeing the stove), or both (American Psychiatric Association, 2013; Mackenzie et al., 1995; Rachman, 2002; Rachman and de Silva, 1978). SIC was predictably triggered by external sensory cues (presentation of food pellets and the availability of the water spout). Currently, there is only one suggested animal model of compulsive checking. The quinpirole sensitization model which requires repeated administration of the D2/D3 agonist to produce a preference to one or two objects/locations (Szechtman et al., 2001; Szechtman et al., 1998). One limitation of this paradigm is the requirement of a pharmacological manipulation and according to diagnostic criteria in humans, the compulsion must not be attributable to the physiological effects of substances or medications (American Psychiatric Association, 2013). We propose SIC as a potentially more naturalistic alternative to study the neurobiology of compulsive checking.

According to Szechtman et al., (Szechtman et al., 1998, Szechtman et al., 2001), the quinpirole sensitization model of checking shares a formal conceptual framework/etiological criterion with compulsive checking observed in OCD that includes, a) exaggerated preoccupation expressed as a physical hesitancy to leave items of interest, b) perseverative, ritualistic motor activity, and c) dependence of checking behaviour on the environmental context. SIC, as *observed* here, also meets this criterion. However, this definition may lack consideration for important cognitive features found in OCD. For instance, a novel rodent compulsive checking model details an operant task(s) examining behaviours that authors postulate underlie the obsessional thought disorders associated with checking in OCD (Eagle et al, 2014; d’Angelo et al, 2017). These models include measures of ‘information gathering’ that mitigate ‘uncertainty’ hypothesized to reduce anxiety. Whether these cognitive components are features of the SIC observed here cannot be determined at this time but deserve future investigation.

### Neural circuits involved in SIP and SIC in rats

Considering the potential role of the oval sub-region of the BNST in energy homeostasis, we expected robust neurophysiological alterations in a paradigm such as SIP, that requires a long-term caloric challenge. From a circuitry perspective, the ovBNST receives its main excitatory inputs from the paraventricular region of the thalamus (PVT) (Li and Kirouac, 2008; Moga et al., 1995). The PVT is important in energy homeostasis, and is involved in anticipatory-feeding behaviours following low caloric intake (de Vasconcelos et al., 2006; Kelley et al., 2005; Nakahara et al., 2004). In return, the ovBNST sends a monosynaptic GABA projection to the lateral hypothalamus (LH) to regulate feeding behaviours (Jennings et al., 2013). The possible role of the LH in encoding learned responses in association with reward associated cues could also influence the establishment of the SIP/SIC action that becomes habitual/compulsive in nature (Nakamura et al., 1987; Nieh et al., 2015). Further to this, the ovBNST’s reciprocal projections to the central amygdala (CeA) may also play a part in transitioning the SIP/SIC behavior from a habit to compulsion as the integrity of the central amygdala function and therefore projection to the dorsal lateral striatum (via the substantia nigra) is critical for the expression of compulsive behaviors (Murray et al, 2015; Everitt and Robbins, 2016). Finally, it is possible that demyelination of key projections within the ovBNST and the above suggested circuits could contribute to and/or explain a predisposition to dysregulated GABA synaptic plasticity in HD rats (Navarro et al, 2017), thus a state we hypothesize could precipitate SIP/SIC behaviours. However, these speculations require evidential substantiation.

The ovBNST is also positioned to exert powerful regulation of the HPA axis through a di-synaptic connection with the paraventricular nucleus of the hypothalamus through the fusiform region of the BNST. Since compulsive behaviours in general, and the SIP paradigm in particular, are linked with chronic stress and pathological anxiety states, it is possible that dysregulated ovBNST function may contribute significantly in these phenomena. Indeed, there is evidence that the ovBNST contributes in adaptive and maladaptive anxiety states (Kim et al., 2013; Normandeau et al., 2018), yet the consequences of altered satiety-state dependent bi-directional GABA plasticity we observed with SIP and SIC on ovBNST input-output neurophysiology is unknown. Furthermore, the behavioural sequelae (i.e., manifestation and expression of SIP/SIC) of aberrant GABA plasticity in the ovBNST within the “compulsion” circuit still has to be revealed.

Further experiments should confirm whether dysregulated GABA plasticity in the ovBNST is the cause or consequence of the behavioural phenomena, and accordingly, either a potential therapeutic target or predictive marker of compulsivity. We also propose that time spent at the water spout could be the primary variable to consider when investigating compulsivity using the SIP rodent model. Future studies should include many behavioural measures of SIP/SIC, including licks at the bottle and/or food magazine entries, as this additional information may shed light on important motor, motivational, or other components of SIP/SIC to further clarify the compulsive nature of these behaviors. Because SIC resisted removal of the caloric challenge and remained adjunctive to food pellet delivery, it could represent a powerful pre-clinical measure of cue-triggered compulsive behaviours.

## Acknowledgements

JGG was funded by NSERC Vanier Graduate Scholarship; SA was funded by a Queen Elizabeth II Graduate Scholarship in Science and Technology; ERH was funded by CIHR Postdoctoral Fellowship (MFE-123712); ÉCD was funded by the Canadian Institute of Health Research (MOP-25953).

## References

American Psychiatric Association, 2013. Diagnostic and statistical manual of mental disorders: DSM-5., Washington, DC.

Belin-Rauscent, A., Daniel, M.L., Puaud, M., Jupp, B., Sawiak, S., Howett, D., McKenzie, C., Caprioli, D., Besson, M., Robbins, T.W., Everitt, B.J., Dalley, J.W., Belin, D., 2016. From impulses to maladaptive actions: the insula is a neurobiological gate for the development of compulsive behavior. Mol Psychiatry 21(4), 491–499.

Berman, I., Kalinowski, A., Berman, S.M., Lengua, J., Green, A.I., 1995. Obsessive and compulsive symptoms in chronic schizophrenia. Compr Psychiatry 36(1), 6–10.

Brett, L.P., Levine, S., 1979. Schedule-induced polydipsia suppresses pituitary-adrenal activity in rats. J Comp Physiol Psychol 93(5), 946–956.

Brett, L.P., Levine, S., 1981. The pituitary-adrenal response to “minimized” schedule-induced drinking. Physiol Behav 26(2), 153–158.

d’Angelo, C., Eagle, D. M., Coman, C. M., & Robbins, T. W. (2017). Role of the medial prefrontal cortex and nucleus accumbens in an operant model of checking behaviour and uncertainty. Brain and neuroscience advances, 1, 2398212817733403.

de Vasconcelos, A.P., Bartol-Munier, I., Feillet, C.A., Gourmelen, S., Pevet, P., Challet, E., 2006. Modifications of local cerebral glucose utilization during circadian food-anticipatory activity. Neuroscience 139(2), 741–748.

Drubach, D.A., 2015. Obsessive-compulsive disorder. Continuum (Minneap Minn) 21(3 Behavioral Neurology and Neuropsychiatry), 783–788.

Eagle, D. M., Noschang, C., d’Angelo, L. S. C., Noble, C. A., Day, J. O., Dongelmans, M. L., … & Robbins, T. W. (2014). The dopamine D2/D3 receptor agonist quinpirole increases checkinglike behaviour in an operant observing response task with uncertain reinforcement: a novel possible model of OCD. Behavioural brain research, 264, 207–229.

Everitt, B.J., Robbins, T.W., 2005. Neural systems of reinforcement for drug addiction: from actions to habits to compulsion. Nat Neurosci 8(11), 1481–1489.

Everitt, B.J., Robbins, T.W., 2016. Drug Addiction: Updating Actions to Habits to Compulsions Ten Years On. Annu Rev Psychol 67, 23–50.

Falk, J.L., 1961. Production of polydipsia in normal rats by an intermittent food schedule. Science 133(3447), 195–196.

Falk, J.L., 1966. The motivational properties of schedule-induced polydipsia. J Exp Anal Behav 9(1), 19–25.

Falk, J.L., 1967. Control of schedule-induced polydipsia: type, size, and spacing of meals. J Exp Anal Behav 10(2), 199–206.

Falk, J.L., 1971. The nature and determinants of adjunctive behavior. Physiol Behav 6(5), 577–588.

Flory, R.K., 1971. The Control of Schedule-Induced Polydipsia: Frequency and Magnitude of Reinforcement. Learn Mem 2, 215–227.

Gregory, J.G., Hawken, E.R., Banasikowski, T.J., Dumont, E.C., Beninger, R.J., 2015. A response strategy predicts acquisition of schedule-induced polydipsia in rats. Prog Neuropsychopharmacol Biol Psychiatry 61, 37–43.

Hariprasad, M.K., Eisinger, R.P., Nadler, I.M., Padmanabhan, C.S., Nidus, B.D., 1980. Hyponatremia in psychogenic polydipsia. Arch Intern Med 140(12), 1639–1642.

Hawken, E.R., Beninger, R.J., 2014. The amphetamine sensitization model of schizophrenia symptoms and its effect on schedule-induced polydipsia in the rat. Psychopharmacology (Berl) 231(9), 2001–2008.

Hawken, E.R., Delva, N.J., Beninger, R.J., 2013a. Increased drinking following social isolation rearing: implications for polydipsia associated with schizophrenia. PLoS One 8(2), e56105.

Hawken, E.R., Delva, N.J., Reynolds, J.N., Beninger, R.J., 2011. Increased schedule-induced polydipsia in the rat following subchronic treatment with MK-801. Schizophr Res 125(1), 93–98.

Hawken, E.R., Lister, J., Winterborn, A.N., Beninger, R.J., 2013b. Spontaneous polydipsia in animals treated subchronically with MK-801. Schizophr Res 143(1), 228–230.

Hooks, M.S., Jones, G.H., Juncos, J.L., Neill, D.B., Justice, J.B., 1994. Individual differences in schedule-induced and conditioned behaviors. Behav Brain Res 60(2), 199–209.

Jennings, J.H., Rizzi, G., Stamatakis, A.M., Ung, R.L., Stuber, G.D., 2013. The inhibitory circuit architecture of the lateral hypothalamus orchestrates feeding. Science 341(6153), 1517–1521.

Kelley, A.E., Baldo, B.A., Pratt, W.E., 2005. A proposed hypothalamic-thalamic-striatal axis for the integration of energy balance, arousal, and food reward. J Comp Neurol 493(1), 72–85.

Kim, S.Y., Adhikari, A., Lee, S.Y., Marshel, J.H., Kim, C.K., Mallory, C.S., Lo, M., Pak, S., Mattis, J., Lim, B.K., Malenka, R.C., Warden, M.R., Neve, R., Tye, K.M., Deisseroth, K., 2013. Diverging neural pathways assemble a behavioural state from separable features in anxiety. Nature 496(7444), 219–223.

Krawczyk, M., Georges, F., Sharma, R., Mason, X., Berthet, A., Bezard, E., Dumont, E.C. Double-dissociation of the catecholaminergic modulation of synaptic transmission in the oval bed nucleus of the stria terminalis. 2011. J Neurophysiol 105(1), 145–153.

Krawczyk, M., Mason, X., DeBacker, J., Sharma, R., Normandeau, C.P., Hawken, E.R., Di Prospero, C., Chiang, C., Martinez, A., Jones, A.A., Doudnikoff, E., Caille, S., Bezard, E., Georges, F., Dumont, E.C., 2013. D1 dopamine receptor-mediated LTP at GABA synapses encodes motivation to self-administer cocaine in rats. J Neurosci 33(29), 11960–11971.

Levine, R., Levine, S., 1989. Role of the pituitary-adrenal hormones in the acquisition of schedule-induced polydipsia. Behav Neurosci 103(3), 621–637.

Li, S., Kirouac, G.J., 2008. Projections from the paraventricular nucleus of the thalamus to the forebrain, with special emphasis on the extended amygdala. J Comp Neurol 506(2), 263–287.

Lockie, S.H., Andrews, Z.B., 2013. The hormonal signature of energy deficit: Increasing the value of food reward. Mol Metab 2(4), 329–336.

López-Crespo, G., Rodríguez, M., Pellón, R., & Flores, P. (2004). Acquisition of schedule-induced polydipsia by rats in proximity to upcoming food delivery. Animal Learning & Behavior, 32(4), 491–499.

Luyten, L., Hendrickx, S., Raymaekers, S., Gabriels, L., Nuttin, B., 2016. Electrical stimulation in the bed nucleus of the stria terminalis alleviates severe obsessive-compulsive disorder. Mol Psychiatry 21(9), 1272–1280.

Mackenzie, T.B., Ristvedt, S.L., Christenson, G.A., Lebow, A.S., Mitchell, J.E., 1995. Identification of cues associated with compulsive, bulimic, and hair-pulling symptoms. J Behav Ther Exp Psychiatry 26(1), 9–16.

Mittleman, G., Jones, G.H., Robbins, T.W., 1988. Effects of diazepam, FG 7142, and RO 15-1788 on schedule-induced polydipsia and the temporal control of behavior. Psychopharmacology (Berl) 94(1), 103–109.

Moga, M.M., Weis, R.P., Moore, R.Y., 1995. Efferent projections of the paraventricular thalamic nucleus in the rat. J Comp Neurol 359(2), 221–238.

Moreno, M., Flores, P., 2012. Schedule-induced polydipsia as a model of compulsive behavior: neuropharmacological and neuroendocrine bases. Psychopharmacology (Berl) 219(2), 647–659.

Murray, J. E., Belin-Rauscent, A., Simon, M., Giuliano, C., Benoit-Marand, M., Everitt, B. J., & Belin, D. (2015). Basolateral and central amygdala differentially recruit and maintain dorsolateral striatum-dependent cocaine-seeking habits. Nature communications, 6, 10088.

Nakahara, K., Fukui, K., Murakami, N., 2004. Involvement of thalamic paraventricular nucleus in the anticipatory reaction under food restriction in the rat. J Vet Med Sci 66(10), 1297–1300.

Nakamura, K., Ono, T., Tamura, R., 1987. Central sites involved in lateral hypothalamus conditioned neural responses to acoustic cues in the rat. J Neurophysiol 58(5), 1123–1148.

Navarro, S. V., Alvarez, R., Colomina, M. T., Sanchez-Santed, F., Flores, P., & Moreno, M. (2016). Behavioral biomarkers of schizophrenia in high drinker rats: a potential Endophenotype of compulsive neuropsychiatric disorders. Schizophrenia bulletin, 43(4), 778–787.

Nieh, E.H., Matthews, G.A., Allsop, S.A., Presbrey, K.N., Leppla, C.A., Wichmann, R., Neve, R., Wildes, C.P., Tye, K.M., 2015. Decoding neural circuits that control compulsive sucrose seeking. Cell 160(3), 528–541.

Normandeau, C.P., Ventura-Silva, A.P., Hawken, E.R., Angelis, S., Sjaarda, C., Liu, X., Pego, J.M., Dumont, E.C., 2018. A Key Role for Neurotensin in Chronic-Stress-Induced Anxiety-Like Behavior in Rats. Neuropsychopharmacology 43(2), 285–293.

Platt, B., Beyer, C.E., Schechter, L.E., Rosenzweig-Lipson, S., 2008. Schedule-induced polydipsia: a rat model of obsessive-compulsive disorder. Curr Protoc Neurosci Chapter 9, Unit 9 27.

Rachman, S., 2002. A cognitive theory of compulsive checking. Behav Res Ther 40(6), 625–639.

Rachman, S., de Silva, P., 1978. Abnormal and normal obsessions. Behav Res Ther 16(4), 233–248.

Rosenzweig-Lipson, S., Sabb, A., Stack, G., Mitchell, P., Lucki, I., Malberg, J.E., Grauer, S., Brennan, J., Cryan, J.F., Sukoff Rizzo, S.J., Dunlop, J., Barrett, J.E., Marquis, K.L., 2007. Antidepressant-like effects of the novel, selective, 5-HT2C receptor agonist WAY-163909 in rodents. Psychopharmacology (Berl) 192(2), 159–170.

Szechtman, H., Ahmari, S.E., Beninger, R.J., Eilam, D., Harvey, B.H., Edemann-Callesen, H., Winter, C., 2017. Obsessive-compulsive disorder: Insights from animal models. Neurosci Biobehav Rev 76(Pt B), 254–279.

Szechtman, H., Eckert, M.J., Wai, S.T., Boersma, J.T., Bonura, C.A., McClelland, J.Z., Culver, K.E., Eilam, D., 2001. Compulsive checking behaviour of quinpirole-sensitized rats as an animal model of Obsessive-Compulsive Disorder (OCD): form and control. BMC Neurosci 2(1), 4.

Szechtman, H., Sulis, W., Eilam, D., 1998. Quinpirole induces compulsive checking behavior in rats: a potential animal model of obsessive-compulsive disorder (OCD). Behav Neurosci 112(6), 1475–1485.

Tazi, A., Dantzer, R., Mormede, P., Le Moal, M., 1986. Pituitary-adrenal correlates of schedule-induced polydipsia and wheel running in rats. Behav Brain Res 19(3), 249–256.

Toscano, C.A., Kameyama, M., Garcia-Mijares, M., Silva, M.T., Santarem, E.M., 2008. Relationship between ethanol and sucrose self-administration and schedule-induced polydipsia. Pharmacol Biochem Behav 90(4), 586–589.

van Kuyck, K., Brak, K., Das, J., Rizopoulos, D., Nuttin, B., 2008. Comparative study of the effects of electrical stimulation in the nucleus accumbens, the mediodorsal thalamic nucleus and the bed nucleus of the stria terminalis in rats with schedule-induced polydipsia. Brain Res 1201, 93–99.

Welkenhuysen, m., Gligorijevic, I., Ameye, L., Prodanov, D., Van Huffel, S., Nuttin, B., 2013. Neuronal activity in the bed nucleus of the stria terminalis in a rat model for obsessive-compulsive disorder. Behav Brain Res 240, 52–59.

